# Suppression by RNA Polymerase I Inhibitors Varies Greatly Between Distinct RNA Polymerase I Transcribed Genes in Malaria Parasites

**DOI:** 10.1101/2024.09.02.610888

**Authors:** Hermela Samuel, Riward Campelo-Morillo, Björn F.C. Kafsack

## Abstract

Transcription of ribosomal RNA (rRNA) by RNA Polymerase I (Pol I) is the rate-limiting step in ribosome biogenesis and a major determinant of cellular growth rates. Unlike virtually every other eukaryote, which express identical rRNA from large tandem arrays of dozens to hundreds of identical rRNA genes in every cell, the genome of the human malaria parasite *Plasmodium falciparum* contains only a handful single-copy 47S rRNA loci that differ substantially from one another in length, sequence and expression in different cell-types. We found that growth of malaria parasite was acutely sensitive to the Pol I inhibitors 9-hydroxyellipticine and BMH-21 and demonstrate that they greatly reduce the transcription of 47S rRNAs as well as transcription of other non-coding RNA genes. Surprisingly, we found that the various types of Pol I-transcribed genes differed by more than two orders of magnitude in their susceptibility to these inhibitors and explore the implications of these findings for regulation of rRNA in *P. falciparum*.

## INTRODUCTION

Ribosomes are macromolecular machines that translate the genetic information within messenger RNA into proteins that carry out the vast majority of biological functions (Petrov et al. 2015; Fox 2010; Cooper 2000). Given the ribosome’s central role in cellular function, eukaryotic genomes encode hundreds of identical ribosomal RNA (rRNA) genes that are organized into large tandem arrays and ribosome composition tends to be highly uniform across cell types (Yang and Karbstein 2024; Goodfellow and Zomerdijk 2013; Sáez-Vásquez and Delseny 2019).

Malaria parasites represent a striking exception to this rule. Despite having similarly sized genomes as budding yeast, which has a single array of 150 identical rRNA copies, the genomes of malaria parasites generally contain four to six rRNA loci, each located on a different chromosome and encoding only a single 47S rRNA gene (McGee, Armache, and Lindner 2023; Rodríguez-Almonacid et al. 2023; Waters et al. 1997; Waters, Syin, and McCutchan 1989). Moreover, this small number of genes encode 2-3 distinct forms of 47 rRNAs that differ substantially from one another in length, sequence and their expression throughout the parasite lifecycle. The major human malaria parasites *Plasmodium falciparum* has 5 complete 47S rRNA genes. Two nearly identical A-type rRNA loci (A1 and A2 on chromosomes 5 and 7) are expressed in the liver and asexual blood stages, the S1 locus on chromosome 1 that is primarily expressed during gametocyte development, and two nearly identical S2-type loci (S2a and S2b on chromosomes 11 and 13) that are highly expressed in the mosquito stages (Waters, Syin, and McCutchan 1989; Rogers et al. 1996; Zhu et al. 1990). Regulation of the S2 loci is also quite distinct in *P. falciparum*. Under standard culture at 37°C, these loci are only weakly expressed but their expression increases rapidly if parasites are exposed to the same temperatures experienced in the mosquito (Fang and McCutchan 2002; Fang, Sullivan, and McCutchan 2004; Sharma et al. 2023).

### 47S transcription machinery

In eukaryotes, transcription is carried out by three RNA polymerase complexes (Pol I, II, and III), that share some subunits but have distinct catalytic cores, promoter types and recruitment machinery. Protein coding genes are transcribed by Pol II to generate capped and polyadenylated messenger RNAs that are targeted to the ribosomes for translation. Pol III transcribes the 5S rRNA, transfer RNAs, and other small non-coding RNAs with a host of functions. In contrast, Pol I only generates single transcript in virtually all eukaryotes: the 47S rRNA that is processed into the 28S, 18S, and 5.8S rRNA that form the catalytic core of the ribosome. Despite this specialization transcription by Pol I comprises up to 60% of the total transcriptional activity of growing cells (Warner, Vilardell, and Sohn 2001). Transcription of the 47S rRNA is the rate limiting step in ribosome biogenesis (Moss and Stefanovsky 1995) and a major determinant of cellular growth rates (Teng, Thomas, and Mercer 2013; Hwang and Denicourt 2024; Kang et al. 2021), and commonly upregulated in a variety of cancer types. Unlike the activity of the other two RNA polymerases, transcription by Pol I is insensitive to inhibition by α-amanitin (Bensaude 2011) but Pol I-specific inhibitors have recently been developed for use in oncotherapy (Laham-Karam et al. 2020).

Since malaria parasites are highly unique in their diversity, organization, and expression of the 47S ribosomal RNA, we decided to explore the effect of these inhibitors on growth and Pol I transcription in the most wide-spread and deadly human malaria parasite *P. falciparum*.

### MATERIALS AND METHODS

### Parasite culture

In this study we used the *Plasmodium falciparum* parasite strain NF54 obtained from BEI Resources. Parasites were maintained following established cultured conditions (Moll et al. 2008) using 0.5% AlbuMAX II (Gibco) supplemented malaria complete media and kept at 37°C under 5% O_2_, 5% CO_2_, 90% N_2._

### *P. falciparum* growth inhibition assay

A fluorescence-based parasite growth inhibition assay was carried out according to previously described protocols with minor modifications (Johnson et al. 2007; Smilkstein et al. 2004). Briefly, ring-stage parasite cultures were adjusted to 0.5% parasitemia and 2% hematocrit before being seeded into a flat-bottom 96-well plate in the presence of increasing drug concentrations at a final volume of 200 µL. Parasites were allowed to grow at 37°C for 72 hours. Upon incubation time, 150 µL from each well was transferred to a black flat-bottom plate, and 100 µL of SYBR Green lysis buffer (20 mM Tris, pH 7.5; 5 mM EDTA, 0.008% wt/vol saponin, 0.08% vol/vol Triton X-100, and 1X SYBR Green [Invitrogen S7563]) was added to each well. The plate was then incubated with gentle agitation for 1 hour, protected from light.

Fluorescence (excitation at 485 nm/emission at 538 nm) was measured using a SpectraMax iD5 plate reader. Fluorescence units (FU) from three technical replicates were normalized to Chloroquine (50 nM) and DMSO-treated controls (0.5% final concentration), averaged, and plotted using GraphPad Prism software. A four-parameter dose-response curve was fitted, and the EC50 was determined. Reported EC50 values were averaged from three independent experiments, with standard error of mean values.

### RNA Polymerase I Inhibition assay

Cultures of 22 hpi ± 3 hpi trophozoites were adjusted to 1.5% parasitemia, equally split, and treated with increasing concentrations of the RNA Pol I inhibitors 9-HE (5,11-dimethyl-6H-pyrido[4,3-b]carbazol-9-ol, monohydrochloride) CAS 52238-35-4 (Cayman Chemical) and BMH-21 (N-[2-(dimethylamino)ethyl]-12-oxo-12H-benzo[g]pyrido[2,1-b]quinazoline-4-carboxamide) CAS 896705-16-1 (Cayman Chemical). Both treated and untreated parasites (with the same volume of DMSO added) were incubated for 3 hours at 26°C. In parallel, untreated control parasites were kept at 37°C. After the incubation period, the parasites were harvested and washed once with 1xPBS before proceeding to RNA isolation.

### RNA isolation, cDNA synthesis and qPCR

Total RNA from saponin-lysed parasites was extracted by TRIzol reagent (Invitrogen) and using the PureLink RNA Mini Kit (Invitrogen) following manufactures instructions. cDNA was synthesized from 500ng of total RNA pretreated with DNAse I (amplification grade, ThermoScientific) using SuperScript III reverse transcriptase (Invitrogen) and random hexamers. Quantitative PCR was performed on the Quant Studio 6 Flex (Thermo Fisher) using iTaq SYBR Green (BioRad) with specific primers for selected targets (Table 1) and normalized to seryl-tRNA synthetase (PF3D7_0717700).

**Table 1.**
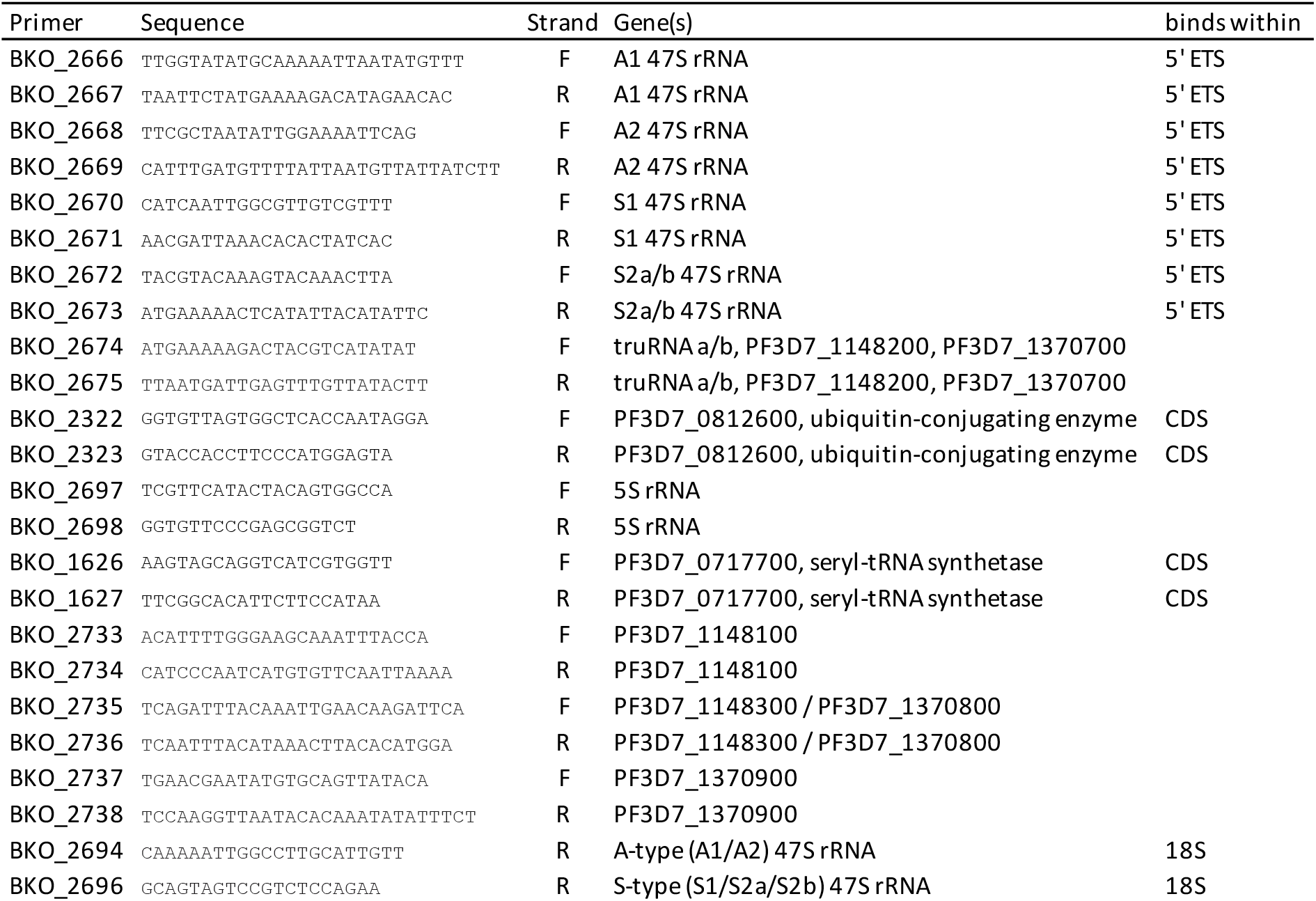
List of primers used in this study.

## RESULTS AND DISCUSSION

### Inhibition of *P. falciparum* RNA polymerase I activity by 9-HE and BMH-21

We decided to test the effect of 9-hydroxyellipticine (9-HE) and BMH-21, two compounds that inhibit RNA Pol I in mammalian cells (Jacobs et al. 2022; Andrews et al. 2013; Kerry et al. 2017). 9-HE was previously shown to have activity against *Plasmodium falciparum* (Montoia et al. 2014) and *Trypanosoma cruzi* (Bénard, Dat-Xuong, and Riou 1975). BMH-21 was shown to inhibit growth and RNA Pol I transcription in African trypanosomes (Kerry et al. 2017) but its anti-malarial activity has not been reported. We found that both compounds were able to inhibit intraerythrocytic replication of *P. falciparum* at sub-micromolar concentrations with EC50 values of 56 ± 3 nM for 9-HE and 352±64 nM BMH-21 (Figure 1).

**Figure 1.**
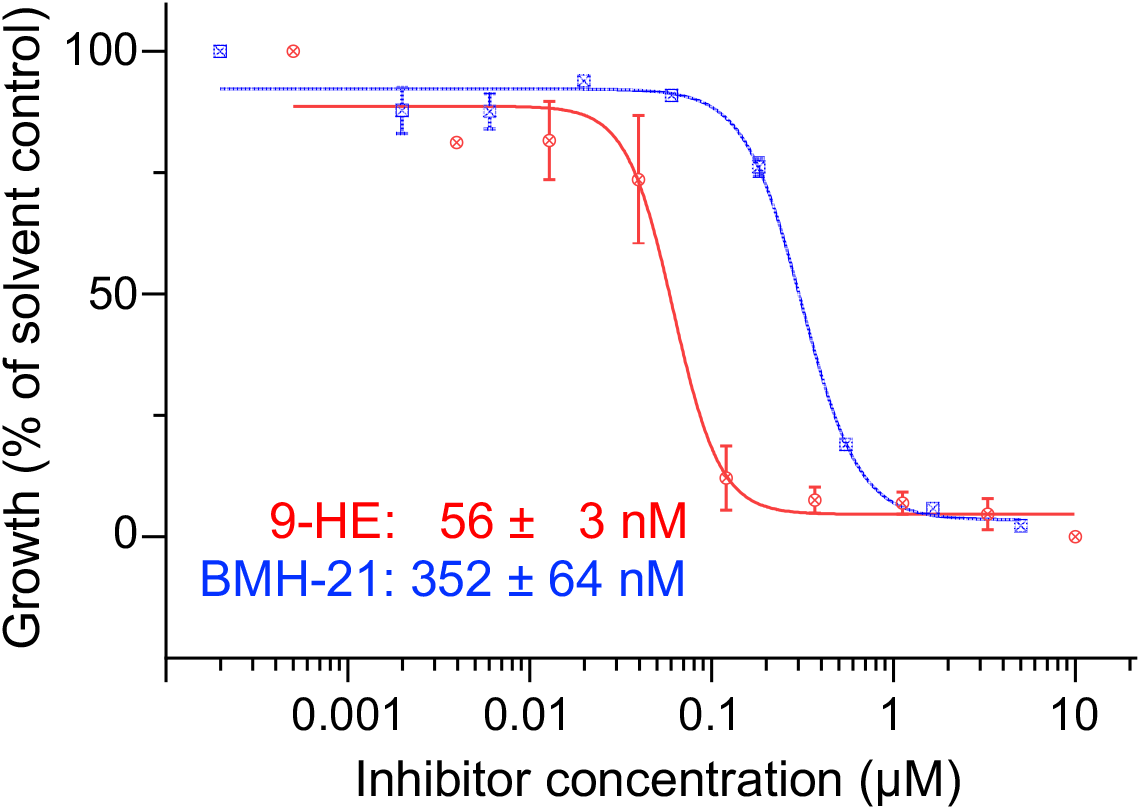
Growth inhibi1on of *P. falciparum* asexual blood stages by 9-HE (red) and BMH-21 (blue). Half maximal effec6ve concentra6on (EC50) ± SEM were calculated from 3 independent biological replicates 72-hour SYBR Green Growth assays. The inhibition curves shown are representative replicates with data points indicating the mean and SEM from 3 technical replicates.

To test if these compounds can inhibit RNA Pol I transcription specifically we monitored transcription of 47S rRNAs, the RNA Pol III-transcribed 5S rRNA, and mRNA transcribed by RNA Pol II. Under standard conditions, only the nearly identical A1 and A2 loci are actively transcribed in asexual blood stages while the S1 and S2a/b loci are expressed in gametocytes and mosquito stages (McGee, Armache, and Lindner 2023). However, transcription of the S2 loci can be induced in asexual blood stage by simply shifting parasite cultures to temperature experienced in the mosquito host thus allowing us to monitor new transcription by RNA Pol I (Fang and McCutchan 2002; Fang, Sullivan, and McCutchan 2004; Sharma et al. 2023). To monitor nascent transcription of 47S rRNA, we used primers the bind within the 5’ External Spacer (5’ETS) that sits upstream of the 18S sequence and is rapidly cleaved and degraded during ribosomal biogenesis (Hughes and Ares 1991; Aubert et al. 2018).

Consistent with these earlier reports (Fang, Sullivan, and McCutchan 2004; Sharma et al. 2023), we found that shifting parasite cultures to 26°C for 3 hours greatly induced expression of S2 47S rRNA and the long non-coding *truRNA* relative to their expression at 37°C (Figure 2A). Transcription of A1 and A2 47S rRNA decreased 2 to 4-fold while transcription of S1 47S rRNA remained largely unchanged. The temperature shift had only minor effects on the 5S and *uce* genes that are transcribed by RNA Pol II and Pol III, respectively. Concurrent treatment with 12.8 µM 9-HE or BMH-21 substantially reduced expression of RNA Pol I transcripts but had no significant effect on the representative Pol II and Pol III transcripts, indicating that both compounds specifically inhibit transcription by RNA Pol I in *P. falciparum* (Figure 2B).

**Figure 2.**
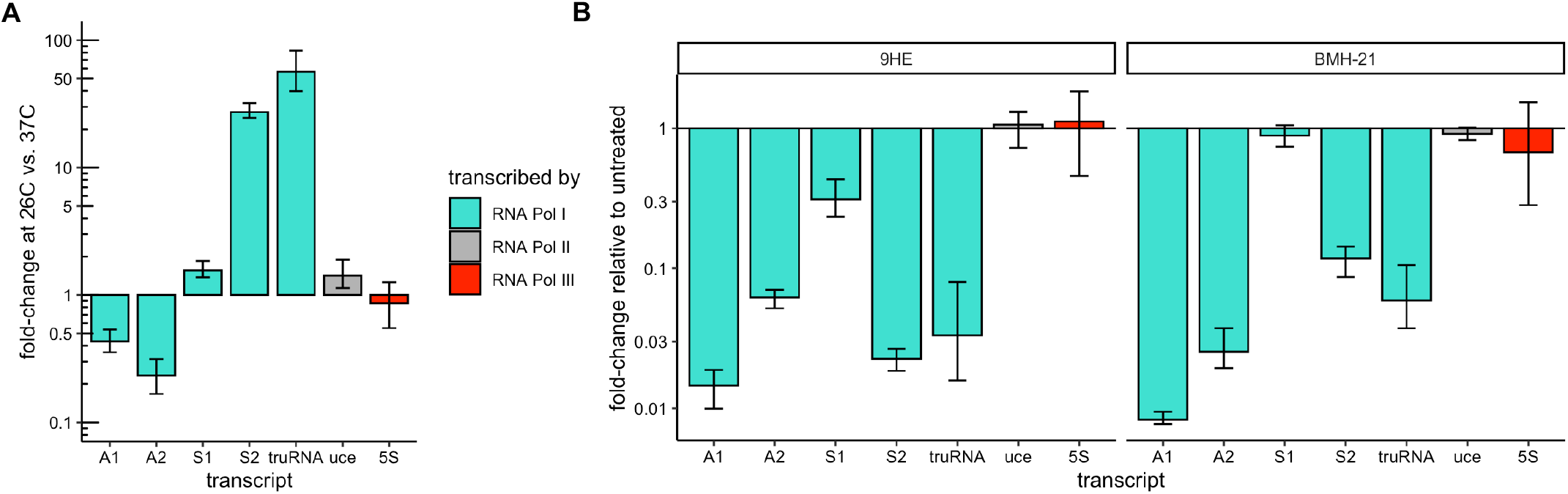
Validation of 9-HE & BMH-21 as specific inhibitors of RNA Polymerase activity in *P. falciparum*. **(A)** Growth at 26°C for 3 hours reduced transcription of A1 and A2 47S rRNA and greatly induces transcription of S2 47S rRNA and the long non-coding *truRNA* but has negligible effects on on S1 47S rRNA, *uce* mRNA, and 5S rRNA transcription. Expression changes of RNA Pol I (turquois), Pol II (grey) and Pol III (red) transcription after 3 hour shift to 26°C relative to expression at 37°C were measured by qRT-PCR. To measure new transcription rather than steady state levels, primers were targeted to the 5’ETS of the 47S rRNAs, which are rapidly cleaved and degraded. **(B)** Concurrent treatment of samples shown in a) with 12.8 µM 9-HE (left panel) or BMH-21 (right panel) inhibits transcription of RNA Pol I transcripts (turquois) but not RNA Pol II (grey) or RNA Pol III (red). The means and non-parametric 95% confidence intervals based on 3 biological replicates are shown throughout.

### Susceptibility to inhibitors of RNA Pol I activity differs greatly between Pol I transcribed genes

Interestingly, we found that transcription of S1 rRNA was less effected by 9-HE and BMH-21 relative to the other four Pol I transcripts (Figure 2B). To test if these other transcripts might also be differentially susceptible to inhibition by either inhibitor, we measured their effect on Pol I transcribed genes across a wider range of concentrations. Indeed, treatment with increasing concentrations of 9-HE or BMH-21 revealed substantially different responses between these RNA Pol I transcribed genes (Figure 3).

**Figure 3.**
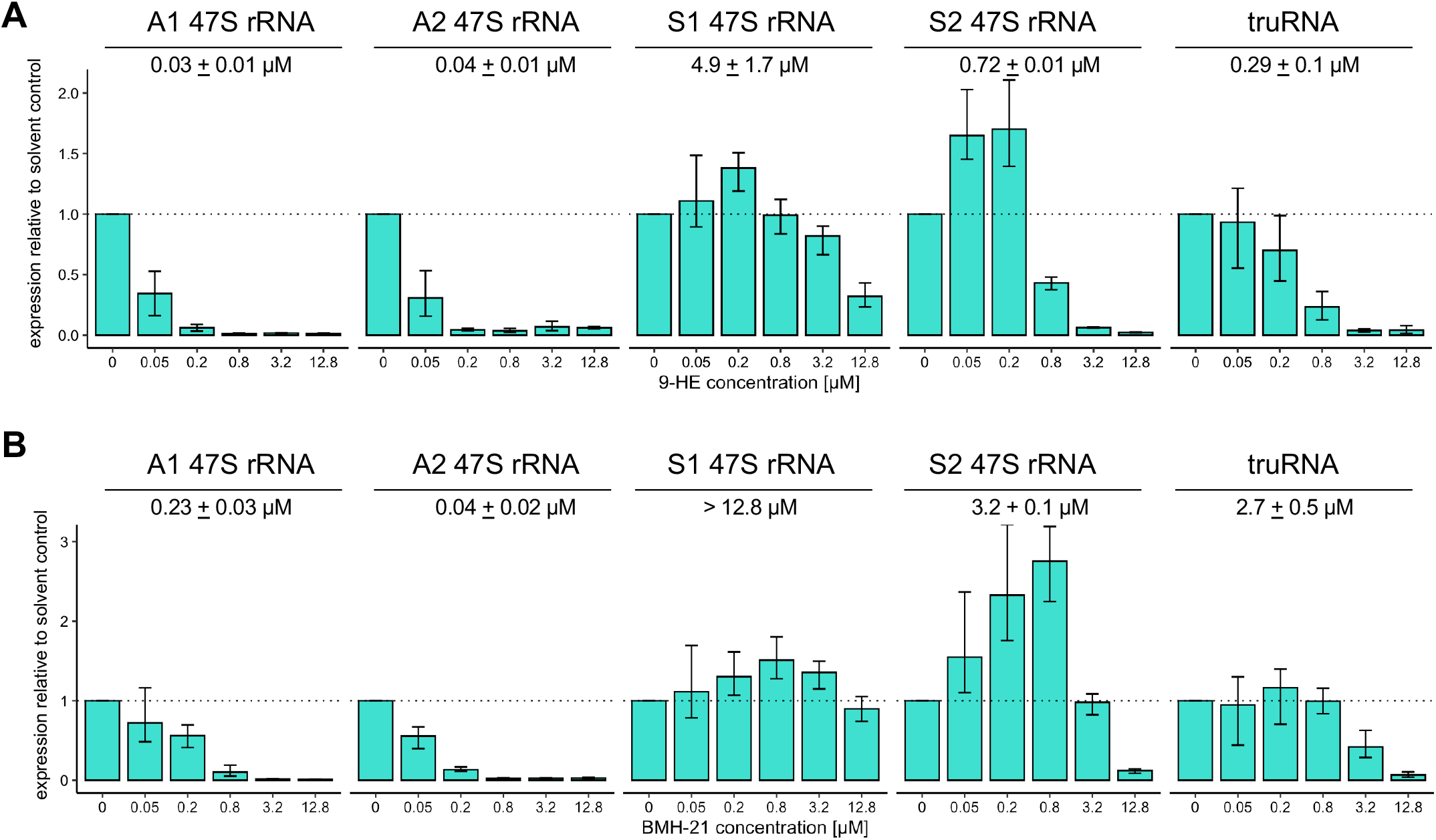
Inhibitor suscep1bility of RNA Pol I transcrip1on differs substan1ally between loci. Transcrip6on of A1, A2, S1, and S2 47S rRNA or *truRNA* in the presence of five different concentra6ons of 9-HE **(A)** or BMH-21 **(B)** was measured by qRT-PCR and compared to solvent control (doTed line). Bars indicate the mean change relative to solvent control, the error bars indicate the 95% confidence interval and the numerical values shown are the mean half-maximal concentra6ons (IC50) based on 3 independent biological replicates.

Transcription of A-type 47S rRNAs was acutely sensitive with half-maximal inhibition concentrations in 30nM range for 9-HE and 140 nM for BMH-21. These concentrations are a close match to those inhibiting growth, strongly suggesting the mechanism of action of 9-HE and BMH-21 is indeed the inhibition of A-type rRNA transcription, which form the vast majority of ribosomes in asexual blood stages. Strikingly, inhibiting the transcription of the other genes required much higher concentrations of 9-HE with half-maximal concentrations of 290 nM, 720 nM, and 4.9 µM for *truRNA*, S2 rRNA, and S1 rRNA respectively. Inhibition of transcription by BMH-21 at these loci followed the same pattern (A > S2 = *truRNA* > S1) but was approximately 4-fold less susceptible compared to 9-HE.

Intriguingly, transcription of S1 and S2 rRNAs increased at lower concentrations that effectively inhibit transcription of A-type rRNAs but not transcription of the *truRNA*. We therefore observed four distinct patterns of susceptibility to these Pol I inhibitors: (i) high susceptibility for A-type rRNA, (ii) intermediate susceptibility with enhanced transcription at concentrations that already inhibit A-type rRNA for S2 rRNA, (iii) intermediate susceptibility without enhancement for the *truRNA*, and (iv) low susceptibility with enhancement for S1 rRNA.

The observation that transcription of these genes differs substantially in their susceptibly to Pol I inhibitors argues for a sequence dependence of this activity, though it could also be explained to differences in the Pol I complex composition that alters inhibitor susceptibility.

Both inhibitors have been shown to intercalate into GC-rich DNA (Elcock, Rodger, and Richards 1996; Jacobs et al. 2022) but with different effects. Treatment with BMH-21 most strongly alters transcription elongation by Pol I and leads to stalling of Pol I upstream of G-rich stretches in the template (Jacobs et al. 2022) while 9-HE specifically altered promoter binding of the SL1 promoter recognition complex required for recruitment of Pol I and transcription initiation (Andrews et al. 2013). Fang and colleagues noted that the transcription start site of A1 and A2 rRNA genes is immediately preceded by a 20nt stretch of 75% GC content that is absent from the S1 and S2a/b promoters (Fang, Sullivan, and McCutchan 2004), providing a likely explanation for the greater inhibitors sensitivity of A-type transcription we observed. Interestingly, the 100bp immediately upstream of the S2a and S2b transcription start sites contains six regularly spaced deoxyguanosine di-nucleotides (14,16,16,17, and 19 bp apart) that stand out within this region of only 18% GC content, while no such evenly spaced repeats could be found close to the *truRNA* or S1 rRNA transcription start sites.

The observed enhancement of S1 and S2a/b transcription at concentrations that inhibited A1/A2 transcription suggests a limited supply of a shared component of the transcriptional machinery that is not targeted by the inhibitors. Reduced transcription at the more susceptible A-type loci at lower temperatures then frees these up for additional transcription of S1 and S2. Intriguingly, the lack of enhancement at the same concentrations would imply that this shared component is not limiting for truRNA transcription. The importance of such repeats for S2 rRNA transcription warrants further study.

### lncRNAs downstream of the *truRNA* likely derive from a single longer transcript

The current *P. falciparum* genome annotations (PlasmoDB release 68) include three non-coding RNA genes (PF3D7_1148100/200/300, referred to as regions *a/b/c* hereafter) immediately upstream of the S2b locus and three additional non-coding RNA genes (PF3D7_1370500/600/700, referred to as regions *d/e/f* hereafter) upstream of the S2a locus. Of these, regions *b-f* lie upstream of the S2a and S2b rRNA genes on chromosomes 13 and 11 within15.4 kb regions that share 98.6% sequence identity (Figure 4A). Regions *a* and *d* are contained within the gene that produces the recently described 2.8 kb *truRNA* (Sharma et al. 2023) and region *c* is homologous to sequences within region *e*.

**Figure 4.**
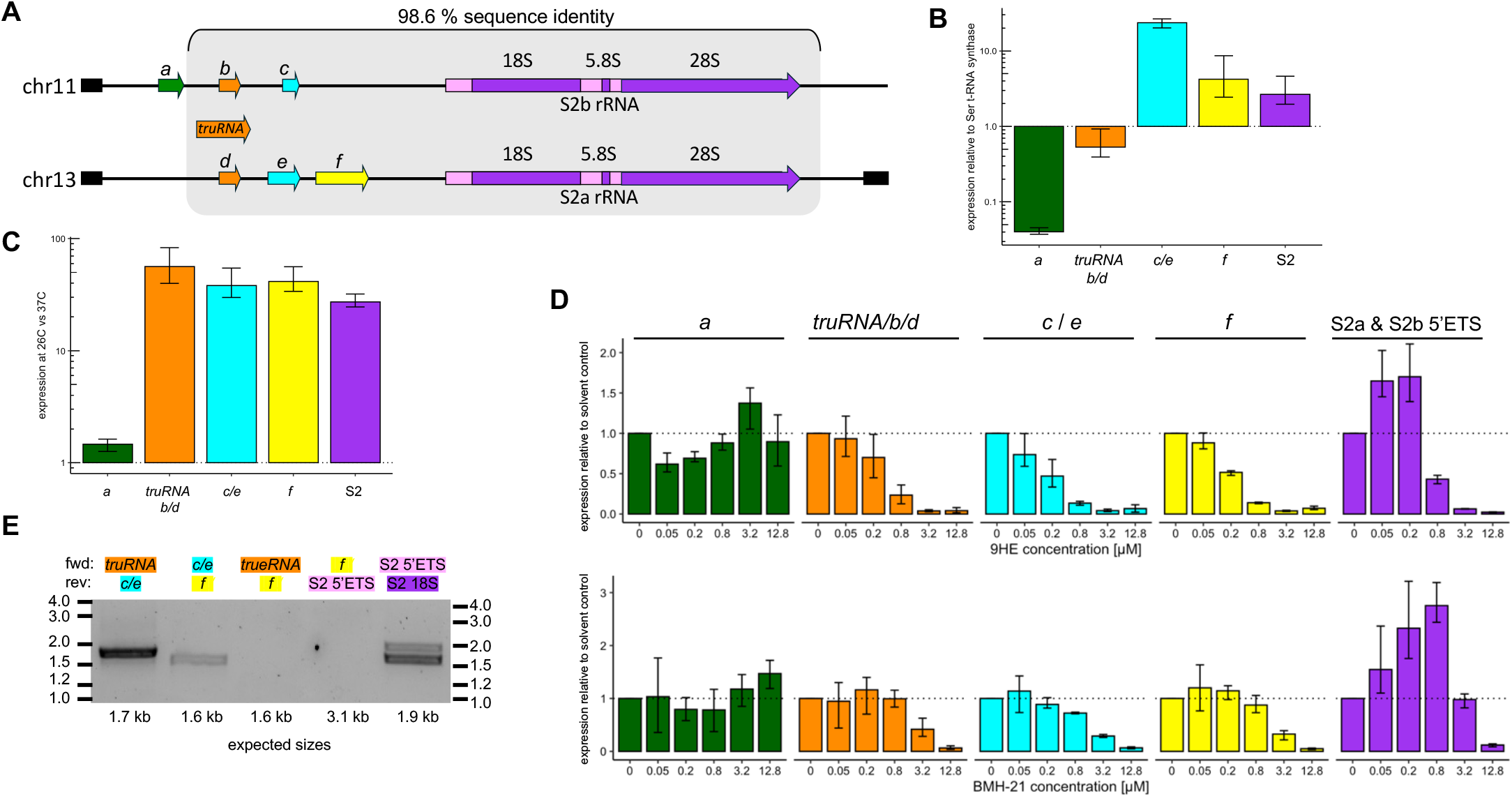
Regulation of S2-locus associated lncRNAs. **(A)** Organization of the S2a and S2b rRNA loci (purple) on chromosomes 13 and 11 along with annotated upstream non-coding RNAs. Six regions (*a/b/c* on chr. 11 and *d/e/f* on chr. 13) are annotated as non-coding RNA genes PF3D7_1148100/200/300 and PF3D7_1370500/600/700, respectively, in the current version of PlasmoDB (release 68). Of these, regions b-f fall within a 15 kb span of very high homology (grey) while region *a* is only found on chromosome 11 immediately upstream. Regions *b* and *d* (orange) are contained within the gene that produces the recently described 2.8 kb truRNA. The nearest protein coding genes are indicated in black. **(B)** Transcript levels relative to seryl-tRNA-synthase. **(C)** Change in expression from regions *a-f* and S2 rRNA after a 3 hour shift to 26°C. **(D)** Transcription from regions *a-f* and S2 rRNA in the presence of five different concentrations of 9-HE (top) and BMH-21 (bottom) was measured by qRT-PCR and compared to solvent control (dotted line). **(E)** RT-PCR products from primers spanning the indicated regions amplifying cDNA from the solvent control sample. Expected size are indicated below each lane and primers targeting the nascent S2 47S rRNA were used a positive control. Amplification from a “no RT” control yielded no products (not shown). All bars indicate the mean and the error bars show the 95% confidence interval based on 3 independent biological replicates.

Since both the S2-type rRNA and the *truRNA* are rapidly upregulated by exposure to lower temperatures, we wanted to test whether the transcripts from regions *a, b/d, c/e*, or *f* might be similarly regulated. We detected substantial transcript being made from all regions (Figure 4A but found that only the regions overlapping with or downstream of the *truRNA* were cold-inducible while expression of PF3D7_1148100 was effectively unchanged (Figure 4B). Transcription of PF3D7_1148100 was also not sensitive to RNA Pol I inhibitors, while the inhibition patters for the other mirrored that of *truRNA*. While regions *c/e* and *f* had distinct steady state levels from the *truRNA* (Figure 4B), the fact that they exhibited highly similar upregulation in response to temperature shift and their susceptibility to RNA Pol I inhibitors lead us to wonder whether they might be transcribed from a single promoter prior to subsequent processing into fragments with variable stability. Indeed, we were able to amplify transcripts bridging from the *truRNA* to region *c/e* and from region *c/e* to region *f* but not from the *truRNA* to region *f* or from region *f* into the *S2a rRNA* (Figure 4E), supporting a single, rapidly processed transcript extending from the *truRNA* into region *f*. The *truRNA* itself was already shown to be processed from a 2.8 kb to a mature 1.3 kb (Sharma et al. 2023). We plan to use long-read RNA sequencing to map the full length transcript in future studies.

When combined with the inability to amplify a product bridging region *f* and the S2 transcript, the distinct inhibitor susceptibility patters of the upstream transcripts and the S2 strongly supports their transcription from distinct promoters (Figure 4D). Current annotations also include a non-coding RNA gene (PF3D7_0531500) immediately upstream of the region encoding the A2 18S rRNA on chromosome 5. However, this annotation should be removed as it is downstream of the A2 rRNA transcription start site and thus part to the A2 5’ETS of the (Fang, Sullivan, and McCutchan 2004).

### *P. falciparum* lacks identifiable homologs to many RNA Pol I-specific components

A search for components of Pol I found in mammals, budding yeast or the protozoan *Trypanosoma brucei*, found only orthologs to RPA1 and RPA2, the two subunits that form the catalytic core (Figure 5). While orthologs to the Pol I subunits shared with Pol II and Pol III were all present, none of Pol I-specific subunits could be identified. Promoter recognition by Pol I depends on distinct complexes in the these other systems (the SL1 complex in humans, CF complex in budding yeast and the CITFA complex in *T*.*brucei*) but again none of the Pol I specific components appear to be present in the malaria parasites. Pol I transcriptional machinery in malaria parasites therefor is either highly divergent or substantially reduced and warrants further investigation.

**Figure 5.**
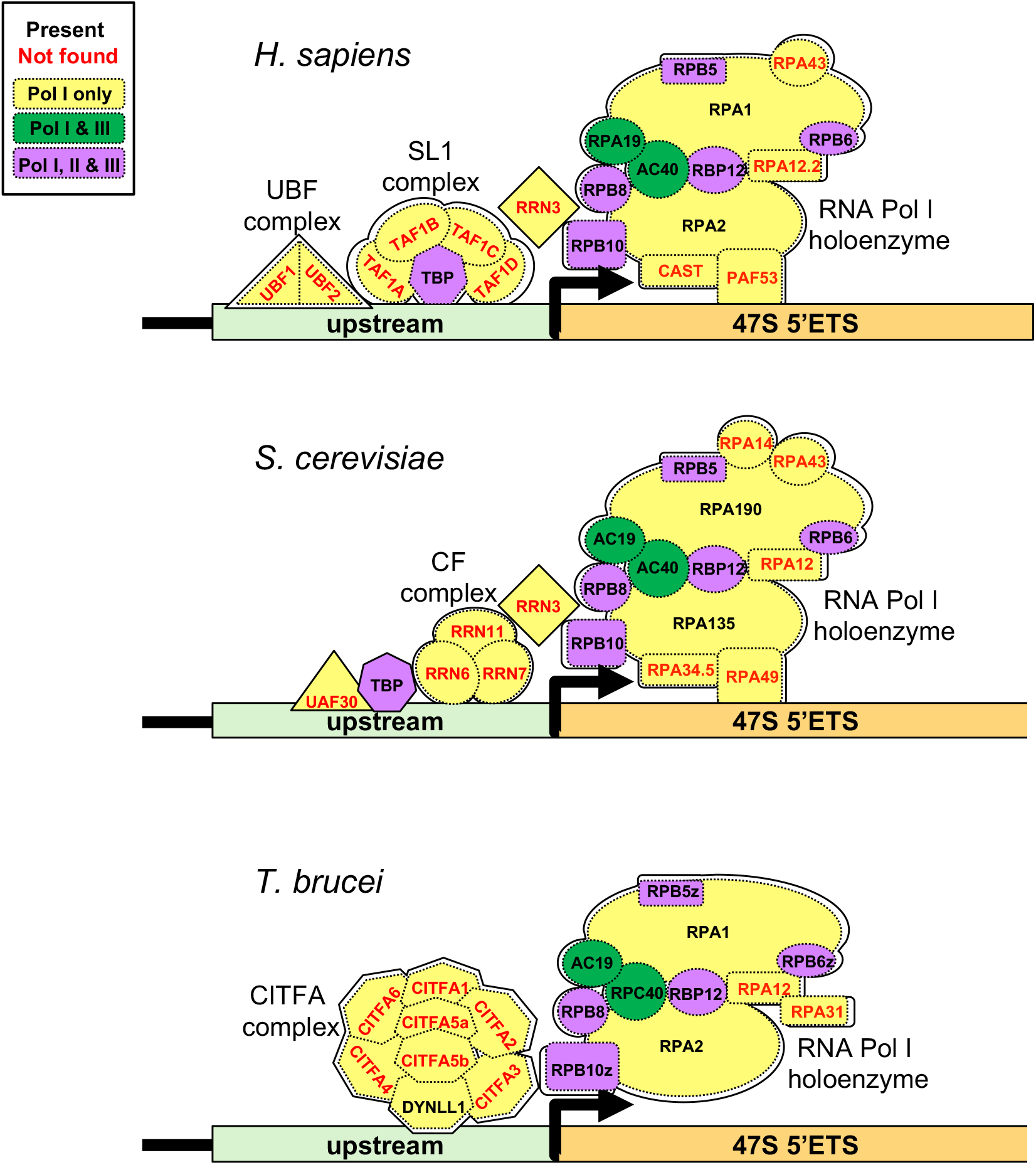
*P. falciparum* lacks identifiable homologs to many RNA Pol I-specific components. Subunits specific to RNA Polymerase I holoenzyme or the promoter recognition complexes are shown in yellow, those shared with RNA Pol III in green, and those shared by RNA Pol I, II and III in purple. Black names indicate subunits with identifiable orthologs in *P. falciparum* while no clear ortholog exists for those with red names.

## Author Contributions

Conceptualization, BK & RC; Methodology, RC; Validation, RC; Formal Analysis, HS & RC.; Investigation, HS & RC; Writing – Original Draft Preparation, RC; Writing – Review & Editing, BK & RC; Visualization, HS, RC, & BK; Supervision, BK & RC; Project Administration, BK.; Funding Acquisition, BK.

## Data Availability Statement

All Data are contained within the manuscript.

## REFERENCES

Andrews, William J., Tatiana Panova, Christophe Normand, Olivier Gadal, Irina G. Tikhonova, and Konstantin I. Panov. 2013. “Old Drug, New Target: ELLIPTICINES SELECTIVELY INHIBIT RNA POLYMERASE I TRANSCRIPTION.” The Journal of Biological Chemistry 288 (7): 4567–82.

Aubert, Maxime, Marie-Françoise O’Donohue, Simon Lebaron, and Pierre-Emmanuel Gleizes. 2018. “Pre-Ribosomal RNA Processing in Human Cells: From Mechanisms to Congenital Diseases.” Biomolecules 8 (4): 123.

Bénard, J., N. Dat-Xuong, and G. Riou. 1975. “Trypanocidal activity of some ellipticine derivatives against Trypanosoma cruzi cultured in vitro.” Comptes rendus hebdomadaires des seances de l’Academie des sciences. Serie D: Sciences naturelles 280 (9): 1177–80.

Bensaude, Olivier. 2011. “Inhibiting Eukaryotic Transcription: Which Compound to Choose? How to Evaluate Its Activity?” Transcription 2 (3): 103–8.

Cooper, GeoWrey M. 2000. Translation of MRNA. Sinauer Associates.

Elcock, A. H., A. Rodger, and W. G. Richards. 1996. “Theoretical Studies of the Intercalation of 9-Hydroxyellipticine in DNA.” Biopolymers 39 (3): 309–26.

Fang, Jun, and Thomas F. McCutchan. 2002. “Thermoregulation in a Parasite’s Life Cycle.” Nature 418 (6899): 742.

Fang, Jun, Margery Sullivan, and Thomas F. McCutchan. 2004. “The EWects of Glucose Concentration on the Reciprocal Regulation of RRNA Promoters in Plasmodium Falciparum.” The Journal of Biological Chemistry 279 (1): 720–25.

Fox, George E. 2010. “Origin and Evolution of the Ribosome.” Cold Spring Harbor Perspectives in Biology 2 (9): a003483.

Goodfellow, Sarah J., and Joost C. B. M. Zomerdijk. 2013. “Basic Mechanisms in RNA Polymerase I Transcription of the Ribosomal RNA Genes.” Sub-Cellular Biochemistry 61: 211–36.

Hughes, J. M., and M. Ares Jr. 1991. “Depletion of U3 Small Nucleolar RNA Inhibits Cleavage in the 5’ External Transcribed Spacer of Yeast Pre-Ribosomal RNA and Impairs Formation of 18S Ribosomal RNA.” The EMBO Journal 10 (13): 4231–39.

Hwang, Sseu-Pei, and Catherine Denicourt. 2024. “The Impact of Ribosome Biogenesis in Cancer: From Proliferation to Metastasis.” NAR Cancer 6 (2): zcae017.

Jacobs, Ruth Q., Abigail K. HuWines, Marikki Laiho, and David A. Schneider. 2022. “The Small-Molecule BMH-21 Directly Inhibits Transcription Elongation and DNA Occupancy of RNA Polymerase I in Vivo and in Vitro.” The Journal of Biological Chemistry 298 (1): 101450.

Johnson, Jacob D., Richard A. Dennull, Lucia Gerena, Miriam Lopez-Sanchez, Norma E. Roncal, and Norman C. Waters. 2007. “Assessment and Continued Validation of the Malaria SYBR Green I-Based Fluorescence Assay for Use in Malaria Drug Screening.” Antimicrobial Agents and Chemotherapy 51 (6): 1926–33.

Kang, Jian, Natalie Brajanovski, Keefe T. Chan, Jiachen Xuan, Richard B. Pearson, and Elaine Sanij. 2021. “Ribosomal Proteins and Human Diseases: Molecular Mechanisms and Targeted Therapy.” Signal Transduction and Targeted Therapy 6 (1): 323.

Kerry, Louise E., Elaine E. Pegg, Donald P. Cameron, James Budzak, Gretchen Poortinga, Katherine M. Hannan, Ross D. Hannan, and Gloria Rudenko. 2017. “Selective Inhibition of RNA Polymerase I Transcription as a Potential Approach to Treat African Trypanosomiasis.” PLoS Neglected Tropical Diseases 11 (3): e0005432.

Laham-Karam, Nihay, Gaspar P. Pinto, Antti Poso, and Piia Kokkonen. 2020. “Transcription and Translation Inhibitors in Cancer Treatment.” Frontiers in Chemistry 8 (April): 276.

McGee, James P., Jean-Paul Armache, and Scott E. Lindner. 2023. “Ribosome Heterogeneity and Specialization of Plasmodium Parasites.” PLoS Pathogens 19 (4): e1011267.

Moll, K., I. Ljungström, H. Perlmann, and A. Scherf. 2008. “Methods in Malaria Research.” Manassas, January. http://kisefront02.ki.se/sites/default/files/methods_in_malaria_research.pdf.

Montoia, Andreia, Luiz F. Rocha E Silva, Zelina E. Torres, David S. Costa, Marycleuma C. Henrique, Emerson S. Lima, Marne C. Vasconcellos, et al. 2014. “Antiplasmodial Activity of Synthetic Ellipticine Derivatives and an Isolated Analog.” Bioorganic & Medicinal Chemistry Letters 24 (12): 2631–34.

Moss, T., and V. Y. Stefanovsky. 1995. “Promotion and Regulation of Ribosomal Transcription in Eukaryotes by RNA Polymerase I.” Progress in Nucleic Acid Research and Molecular Biology 50: 25–66.

Petrov, Anton S., Burak Gulen, Ashlyn M. Norris, Nicholas A. Kovacs, Chad R. Bernier, Kathryn A. Lanier, George E. Fox, et al. 2015. “History of the Ribosome and the Origin of Translation.” Proceedings of the National Academy of Sciences of the United States of America 112 (50): 15396–401.

Rodríguez-Almonacid Cristian Camilos, Morgana K. Kellogg, Andrey L. Karamyshev, and Zemfira N. Karamysheva. 2023. “Ribosome Specialization in Protozoa Parasites.” International Journal of Molecular Sciences 24 (8). 10.3390/ijms24087484.

Rogers, M. J., R. R. Gutell, S. H. Damberger, J. Li, G. A. McConkey, A. P. Waters, and T. F. McCutchan. 1996. “Structural Features of the Large Subunit RRNA Expressed in Plasmodium Falciparum Sporozoites That Distinguish It from the Asexually Expressed Subunit RRNA.” RNA 2 (2): 134–45.

Sáez-Vásquez, Julio, and Michel Delseny. 2019. “Ribosome Biogenesis in Plants: From Functional 45S Ribosomal DNA Organization to Ribosome Assembly Factors.” The Plant Cell 31 (9): 1945–67.

Sharma, Indu, Jun Fang, Eric A. Lewallen, Kirk W. Deitsch, and Thomas F. McCutchan. 2023. “Identification of a Long Noncoding RNA Required for Temperature Induced Expression of Stage-Specific RRNA in Malaria Parasites.” Gene 877 (August): 147516.

Smilkstein, Martin, Nongluk Sriwilaijaroen, Jane Xu Kelly, Prapon Wilairat, and Michael Riscoe. 2004. “Simple and Inexpensive Fluorescence-Based Technique for High-Throughput Antimalarial Drug Screening.” Antimicrobial Agents and Chemotherapy 48 (5): 1803–6.

Teng, Teng, George Thomas, and Carol A. Mercer. 2013. “Growth Control and Ribosomopathies.” Current Opinion in Genetics & Development 23 (1): 63–71.

Warner, J. R., J. Vilardell, and J. H. Sohn. 2001. “Economics of Ribosome Biosynthesis.” Cold Spring Harbor Symposia on Quantitative Biology 66 (0): 567–74.

Waters, A. P., R. M. van Spaendonk, J. Ramesar, R. A. Vervenne, R. W. Dirks, J. Thompson, and C. J. Janse. 1997. “Species-Specific Regulation and Switching of Transcription between Stage-Specific Ribosomal RNA Genes in Plasmodium Berghei.” The Journal of Biological Chemistry 272 (6): 3583– 89.

Waters, A. P., C. Syin, and T. F. McCutchan. 1989. “Developmental Regulation of Stage-Specific Ribosome Populations in Plasmodium.” Nature 342 (6248): 438–40.

Yang, Yoon-Mo, and Katrin Karbstein. 2024. “Ribosome Assembly and Repair.” Annual Review of Cell and Developmental Biology, May. 10.1146/annurev-cellbio-111822-113326.

Zhu, J. D., A. P. Waters, A. Appiah, T. F. McCutchan, A. A. Lal, and M. R. Hollingdale. 1990. “Stage-Specific Ribosomal RNA Expression Switches during Sporozoite Invasion of Hepatocytes.” The Journal of Biological Chemistry 265 (21): 12740–44.

